# Ligand-induced motions in pentameric ligand-gated ion channels revealed by EPR spectroscopy

**DOI:** 10.1101/2020.11.04.368233

**Authors:** Varun Tiwari, Jennifer Borchardt, Abby Schuh, Candice S. Klug, Cynthia Czajkowski

**Author notes:** **Funding:** NIH 34727 to CC, NIH RR022422 and OD025260 to CSK.

## Abstract

Signaling in the brain depends on rapid opening and closing of pentameric ligand-gated ion channels (pLGICs). These proteins are the targets of various clinical drugs and, defects in their function is linked to a variety of diseases including myasthenia, epilepsy and sleep-disorders. While recent high-resolution structures of prokaryotic and eukaryotic pLGICs have shed light on the molecular architecture of these proteins, describing their conformational dynamics in physiological lipids is essential for understanding their function. Here, we used site-directed spin labeling electron paramagnetic resonance (SDSL EPR) spectroscopy and functional channels reconstituted in liposomes to reveal ligand-induced structural changes in the extracellular domain (ECD) of GLIC. Proton-activation caused an inward motion of labeled sites at the top of β-strands (β1, 2, 5, 6, 8) towards the channel lumen, consistent with an agonist-induced inward tilting motion of the ECD. Similar proton-dependent GLIC ECD motions were detected in the presence of a non-activating (gating deficient) mutation, suggesting that the inward tilting of the ECD does not accompany channel opening but is associated with an agonist-induced closed pre-activated channel state. These findings provide new insights into the protein dynamics underlying pLGIC gating transitions.

## Introduction

Pentameric ligand gated ion channels (pLGICs) are allosteric membrane proteins that transduce chemical signals into changes in membrane current. They play important physiological roles in the brain and periphery by mediating fast chemical signaling between cells (Miller and Smart 2010, Corringer, Poitevin et al. 2012). pLGICs carry out their functions by interconverting between three major functional states: resting-closed (unliganded, channel closed), open (liganded, channel open) and desensitized (liganded, channel closed). These global allosteric interconversions are regulated by the binding of ligands, including neurotransmitter, which promote transitions from the resting-closed state to the conducting open state, and eventually, on prolonged occupancy, into a desensitized state (Castillo and Katz 1957, Katz and Thesleff 1957, Corringer, Poitevin et al. 2012). Short lived intermediate ‘pre-active’ closed channel states have also been identified by single channel electrophysiology (Lape, Colquhoun et al. 2008, Mukhtasimova, Lee et al. 2009) and by molecular dynamic simulations(Guros, Balijepalli et al. 2020). The neurotransmitter binding site is contained within the extracellular domain (ECD) of the receptor, but the conformational rearrangements induced by its occupancy extend all the way through the protein to the ion channel in the transmembrane domain (TMD) around 60 Å away (Czajkowski 2005, Miller and Smart 2010, Taly, Hénin et al. 2014, Cecchini and Changeux 2015). How ligand binding promotes changes in structure is one of the central questions about the function of these allosteric proteins.

pLGICs have a conserved 3D structure (Corringer, Poitevin et al. 2012, Taly, Hénin et al. 2014, Cecchini and Changeux 2015, Nemecz, Prevost et al. 2016, Gielen and Corringer 2018). Each pLGIC subunit consists of two parts: the ECD comprises 10 β-strands folded into an immunoglobin-like β-sandwich; the TMD comprises four membrane-spanning α-helices M1-M4. M2 helices from all five subunits come together around a five-fold (or pseudo-five-fold in heteropentamers) symmetry axis to form the central ion pore. The neurotransmitter-binding site is located at the interface between two subunits in the ECD. Flexible loops from each subunit, namely Loop 2, Loop 7, Loop 9 and the M2-M3 loop, connect the ECD to the TMD. Comparing presumed closed and open state structures of the α1 glycine receptor, glutamate-activated chloride channel (GluCl), and *Gloeobacter violaceus* ion channel (GLIC) predicts a common channel opening mechanism involving an outward radial tilting of the extracellular ends of the M2 helix in the TMD and a counterclockwise twisting/inward tilting of the β-sandwich in the ECD (Bocquet, Nury et al. 2009, Hilf and Dutzler 2009, Hibbs and Gouaux 2011, Althoff, Hibbs et al. 2014, Sauguet, Shahsavar et al. 2014, Du, Lu et al. 2015, Nemecz, Prevost et al. 2016). High-resolution structures of the α4β2 nicotinic receptor and the more recent α1β2γ2-GABAA receptor in presumed desensitized states suggest a constriction at the intracellular ends of the pore-lining M2 helix closes the channel during desensitization (Miller and Aricescu 2014, Morales-Perez, Noviello et al. 2016, Laverty, Desai et al. 2019, Masiulis, Desai et al. 2019, Kim, Gharpure et al. 2020). ^19^F NMR experiments on ELIC and mutagenesis studies on GABA_A_ and glycine receptors also support this idea (Gielen, Thomas et al. 2015, Kinde, Chen et al. 2015).

While X-ray crystallography and cryo-electron microscopy have provided detailed structures of multiple pLGICs in different conformations (Amundarain, Ribeiro et al. 2019), their assignment to specific functional states is difficult and molecular mechanisms underlying their allosteric gating transitions remain unclear. Some structures were determined in detergent micelles rather than in lipid bilayers, raising questions about their physiological relevance (Gonzalez-Gutierrez and Grosman 2010, daCosta and Baenziger 2013). Crystal packing and the addition of stabilizing proteins (e.g. nanobodies) to aid in structural determination may preclude capture of key physiological states. The structures are static snapshots. Thus, complementary methods using functional protein are needed to ascertain the dynamics underlying pLGIC gating transitions.

Here, we used site-directed spin labeling electron paramagnetic resonance (SDSL EPR) spectroscopy and functional GLIC channels reconstituted into liposomes to probe motions of the inner β-sheet of the ECD during agonist-induced conformational transitions from the resting-closed state to the desensitized state. SDSL EPR spectroscopy is a powerful biophysical technique that can be used to study structural motions and ligand-induced conformation changes in membrane proteins in their native lipid environment (Hubbell, Cafiso et al. 2000, Klug and Feix 2008, Claxton, Kazmier et al. 2015). This method involves the mutation of targeted residues into cysteine, and their subsequent covalent modification by a sulfhydryl-specific spin label containing an unpaired electron (typically 1-oxyl-2,2,5,5-tetramethyl-Δ3-pyrroline-3-methyl methanethiosulfonate, MTSL). Continuous wave EPR (CW EPR) can then be used to investigate the mobility and local environment of the probe, and double electron-electron resonance (DEER) spectroscopy can measure distances between pairs of spin labels. SDSL EPR spectroscopy is an ideal complement to the static structures determined with X-ray crystallography and electron microscopy, and has been used to study structural dynamics in a number of membrane proteins including rhodopsin, LeuT, β_2_-adrenergic receptor, K^+^-channels, AMPA and NMDA receptors, as well as the pLGIC GLIC (Perozo, Cortes et al. 1998, Altenbach, Kusnetzow et al. 2008, Dellisanti, Ghosh et al. 2013, Dürr, Chen et al. 2014, Kazmier, Sharma et al. 2014, Velisetty, Chalamalasetti et al. 2014, Manglik, Kim et al. 2015, Zhu, Stein et al. 2016). The prokaryotic protein GLIC is homologous with the eukaryotic pLGICs and serves as a useful model for structure-function studies of synaptic receptors. GLIC is especially well suited for SDSL EPR experiments because considerable quantities of recombinant protein can be purified and reconstituted into liposomes, and it has been crystallized in multiple conformations (closed, open, locally-closed and desensitized) (Bocquet, Nury et al. 2009, Hilf and Dutzler 2009, Prevost, Sauguet et al. 2012, Sauguet, Shahsavar et al. 2014). Here, we probed the dynamics of the extracellular ends of β-strands 5, 6, 2, 1, 8 and the β2-β3 loop in the ECD of GLIC. We found that these sites move inwards towards the channel axis during proton-induced transitions from the resting-closed state to the desensitized state, consistent with an agonist-induced inward tilting motion of the ECD inner β-sheet. Similar proton-dependent GLIC ECD motions were detected in the presence of a non-activating (gating deficient) mutation, suggesting that the inward tilting of the ECD does not accompany channel opening but is associated with an agonist-induced closed channel state.

## RESULTS

### Functional characterization of spin-labeled GLIC mutants

Thirteen residues in the ECD on β-strands 5, 6, 2, 1 and 8 (inner β-sheet) were individually mutated into cysteine on a Cys-less GLIC (GLIC C26A) background (**Fig. 1A, B**). GLIC channels were expressed in *E. coli* C43 cells, purified in DDM (n-dodecyl β-D-maltopyranoside) and spin-labeled at the introduced cysteine with MTSL (**Fig. 1C,** see Methods) to form the R1 side chain. Gel filtration chromatography of the spin-labeled proteins showed a clean monodisperse sample eluting at the size of pentameric GLIC (~182 kDa) (**Fig. S1**). To test whether the purified and spin-labeled GLIC proteins were functional, we prepared GLIC proteoliposomes (see Methods) and injected them into *Xenopus* oocytes (Morales, Aleu et al. 1995, Jarecki, Makino et al. 2013). These oocytes had significantly larger proton-induced currents (300-800 nA) than non-injected oocytes (100 nA) demonstrating that each of the spin-labeled GLIC mutants was functional and proton-sensitive (**Fig. 2**).

**Figure 1.**
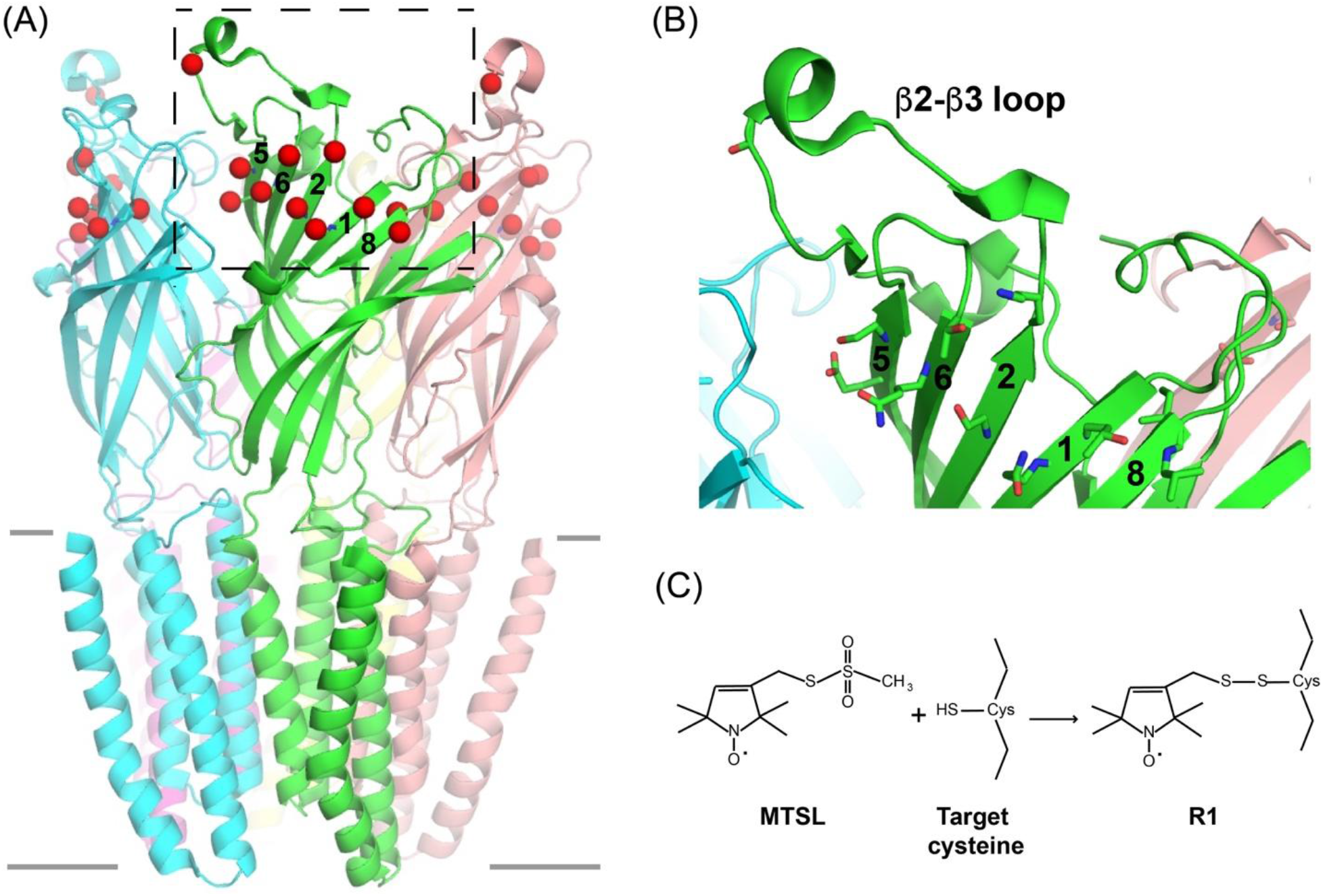
Residues in ECD of GLIC probed using EPR spectroscopy. (A) Side view of GLIC crystal structure (PDB 4HFI) with C_β_ positions of target residues in the inner β-sheet of the ECD shown as red spheres (two subunits in back are hidden for clarity). β-strands in one subunit are numbered. Solid gray lines mark approximate position of lipid bilayer. (B) Expanded view of one subunit highlighting location of spin labeled residues on ends of β5, β6, β2, β1, β8 and in β2-β3 loop as sticks. (C) Scheme showing covalent reaction of MTSL with cysteine to form the R1 side chain.

**Figure 2.**
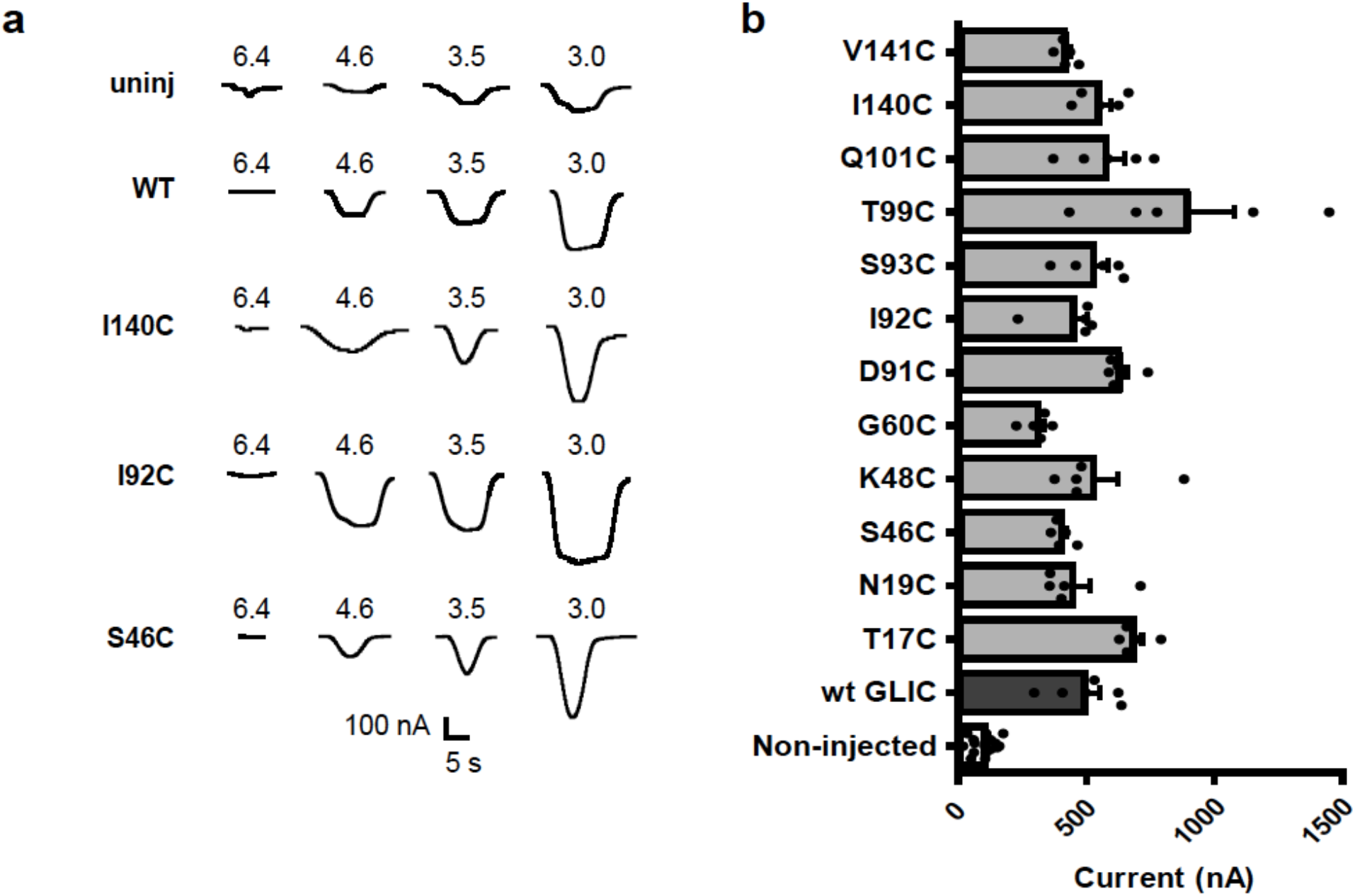
Purified GLIC mutant protein reconstituted into liposomes is functional. (A) Currents elicited by pH jumps measured using two-electrode voltage clamp from non-injected *Xenopus laevis* oocytes and oocytes injected with wild type and spin labeled purified GLIC protein reconstituted into PE:PG liposomes. (B) Summary of pH 3.0 elicited currents from non-injected and oocytes injected with wild type and spin-labeled GLIC proteoliposomes. Bars represent mean +/− SEM from ≥ 5 oocytes. Currents from *Xenopus* oocytes injected with GLIC proteoliposomes were significantly larger than those from non-injected oocytes.

### CW EPR spectroscopy reveals proton-induced changes in spin probe mobility

Spin labeled GLIC was examined by continuous wave (CW) EPR to assess probe mobility and the local packing environment. The shape of a CW spectrum reflects the mobility of the R1 side chain. CW EPR spectra broaden as the motion of the spin label becomes less mobile. To determine which locations in the ECD undergo proton-induced structural rearrangements, we measured room temperature CW spectra for each of the single-cysteine mutants labeled with MTSL at pH 7.6, when the channels were predominantly in a resting-closed state, and at pH 3.0, when the channels were stabilized in a desensitized state. Except for I92C and V94C, the GLIC mutants all had strong CW EPR signals, indicating that the pentamers were modified with two or three spin labels and that the probes were water-accessible. The mobilities of the introduced spin probes depended on their location in the GLIC ECD (**Fig. 3**). The G60R1, V141R1, and S46R1 side chains on GLIC reconstituted into liposomes at pH 7.6 had fast motion (sharper, narrower peaks), indicating that the probes rotate freely in a somewhat unrestricted environment, whereas T99R1, T17R1, N19R1, Q101R1, D91R1, and S93R1, had intermediate motion (broader peaks) suggesting there may be tertiary contacts with neighboring residues, and the spectra of K48R1 and I140R1 indicated very restricted motion.

**Figure 3.**
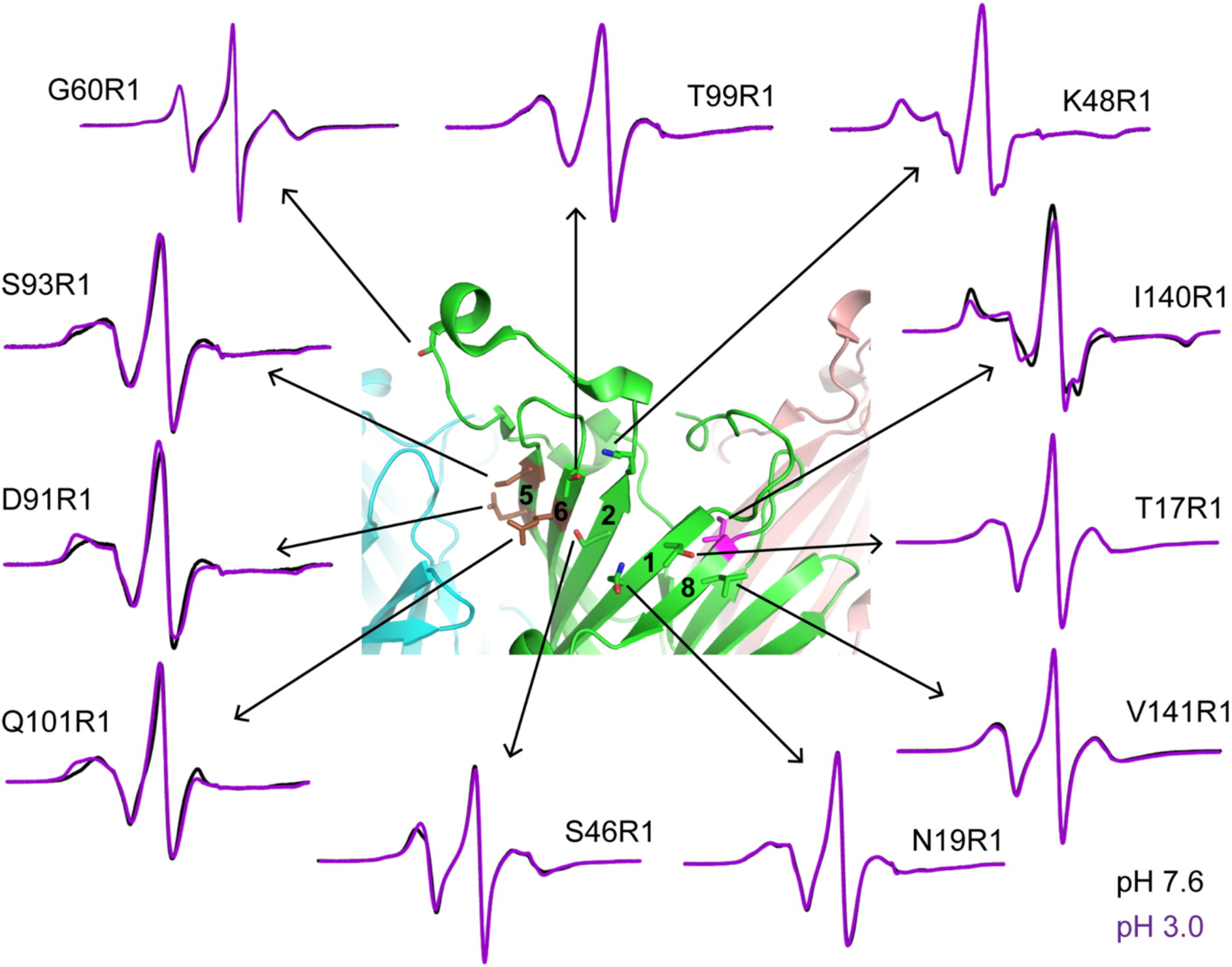
CW EPR spectra of target residues on the inner β-sheet of GLIC ECD. Overlays of X-band CW EPR spectra of spin-labeled protein at pH 7.6 (black, closed state) and pH 3.0 (magenta, desensitized state). Middle panel: Close up view of GLIC crystal structure (4HFI) with spin-labeled positions shown as sticks. R1s that show a pH-induced decrease in spin probe mobility are highlighted in brown, while the R1 that becomes more mobile at pH 3 is highlighted in pink. See **Figure S2** for expanded views of low-field regions highlighting proton-induced changes.

CW spectra were nearly indistinguishable between pH 3.0 and 7.6 for G60R1 (β2-β3 loop), T99R1 (β6 top), K48R1 and S46R1 (β2), T17R1and N19R1 (β1) and V141R1 (β8), indicating the GLIC closed (pH 7.6) to desensitized (pH 3.0) transitions did not alter the packing around these sites (**Fig. 3**). The CW spectra of D91R1, S93R1 (β5) and Q101R1 (β6) became broader at pH 3.0 indicating a proton-induced decrease in mobility (**Fig. 3**, **Fig. S2**), and that the distal ends of β5 and β6 are in a more restricted environment in the desensitized state than in the resting state. In contrast, the CW spectrum of I140R1 (β8), which is pointed toward the inside of the β-sheet, appeared to have two distinct spectral components, mobile and immobile, likely associated with two alternative rotameric spin-probe conformations (**Fig. 3**, **Fig. S2**). At pH 7.6 (resting state), a greater proportion of the spin probe was in an immobile conformation, whereas at pH 3.0 (desensitized state), some of the immobilized spins moved to the more unrestricted environment, indicating a proton-induced structural rearrangement that made the environment near the spin probe on the inside of the sheet somewhat less restricted. Notably, the residues experiencing a decrease in motion upon proton-induced conformational changes are clustered together in strands 5 and 6 and the site exhibiting less restricted mobility is located several strands away on the opposite face of the beta-sheet.

### DEER spectroscopy reveals proton-induced distance changes

DEER spectroscopy reveals distances and distance distributions in the range of 20-60 Å between pairs of spin labels (Klug and Feix 2008, Jeschke 2012, Claxton, Kazmier et al. 2015). We used DEER to measure the distances between probes in the ECD of GLIC at pH 7.6 (resting state) and pH 3.0 (desensitized state) to test for movements of inner β-sheet of the ECD during proton-dependent gating transitions. Comparing GLIC crystal structures in presumed closed (4NPQ, pH 7) and open states (4HFI, pH 4) predicts that channel opening involves an inward tilting motion of the ECD. Whether GLIC ECD motions are associated with transitions into a desensitized state is not well established. **Figure 4D** shows the background-subtracted time-domain DEER signals for T99R1 (β6), fitted using a model-free approach. At pH 7.6 (resting state), the interspin distance distributions for T99R1 on adjacent subunits had a broad peak centered at 46 Å and a slightly narrower peak centered at 42 Å at pH 3.0 (desensitized state) (**Fig. 4E**), indicating that the mean adjacent interspin distance decreased by 4 Å and the spin probes attached to T99C moved closer together at pH 3.0. The widths of the distance distributions primarily reflect the heterogeneity of spin-label orientations at a given site due to multiple MTSL rotamers. Since GLIC is a homopentamer, we expect two distances for each site: one between probes on adjacent subunits and the other between probes on non-adjacent subunits (**Fig. 4B, C**), with a distance ratio of 1.6. Since the non-adjacent distances between probes were beyond the range for accurate measurement using DEER spectroscopy (> 60 Å), they are not reported.

**Figure 4.**
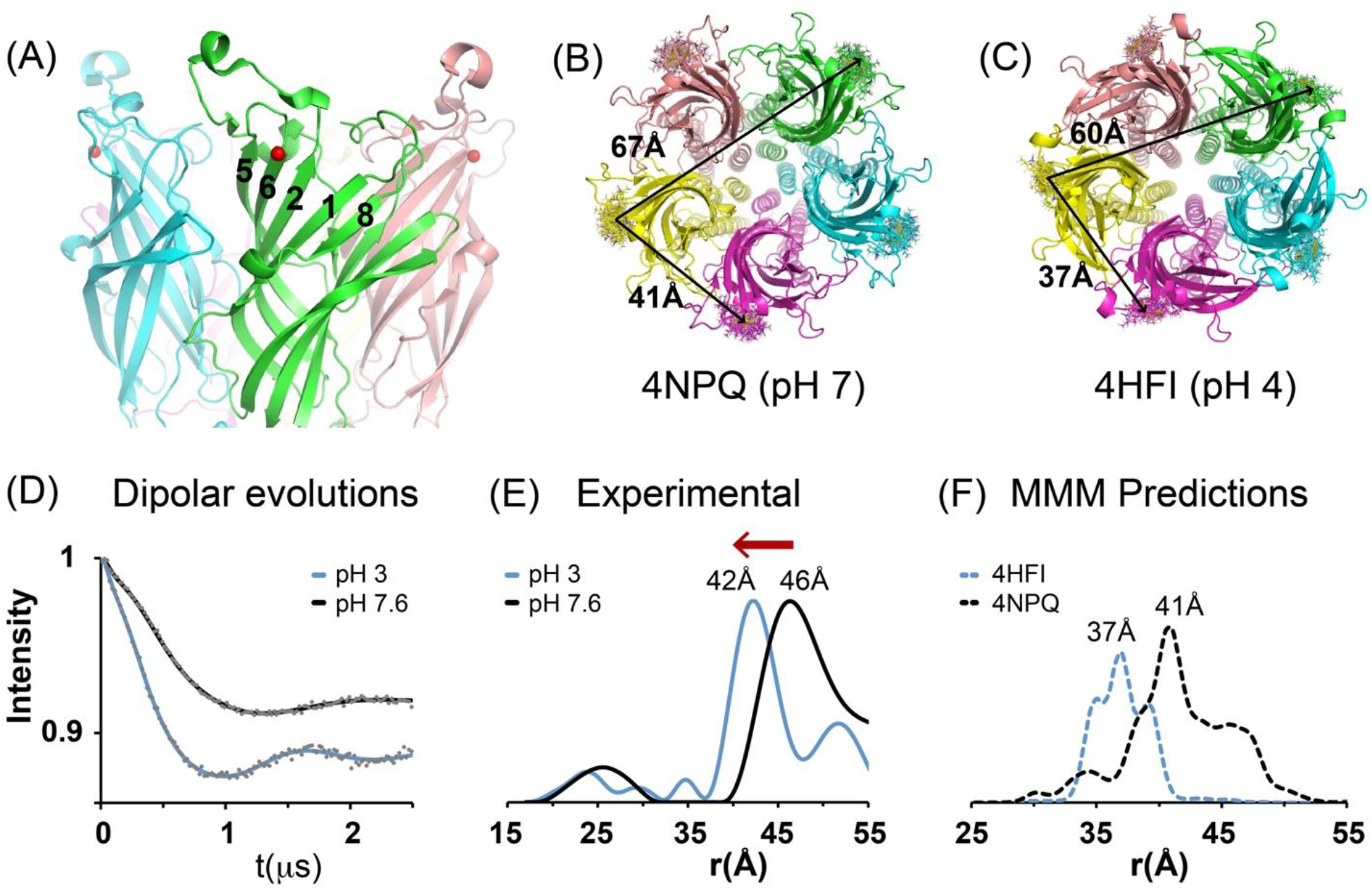
Spin labels attached to T99C (β6) move closer at pH 3.0. (A) Side view of GLIC ECD with C_β_ position of T99 highlighted as a red sphere (B, C) Top-down views of crystal structures of GLIC (4NPQ, pH 7; 4HFI, pH 4) showing the predicted MTSL rotamers (sticks) with black lines depicting the average interspin distances between adjacent & non-adjacent subunits modeled using MMM software. (D) Background subtracted Q-band DEER dipolar evolution data (gray dots) of T99R1 fit using model-free fitting algorithms (pH 7.6, closed state, black solid lines; pH 3.0, desensitized state, blue solid lines) with the program LongDistances (Toledo Warshaviak, Khramtsov et al. 2013). (E) Corresponding experimental interspin distance distributions showing the probability of a distance (P(r)) versus distance (r) are plotted at pH 7.6 (black) & pH 3.0 (blue). Mean distances for highest peaks have been labeled. (F) MMM simulated interspin distance distributions showing the probability of a distance (P(r)) versus distance (r) based on rotamer analysis of T99R1 using crystal structures of GLIC (4NPQ, pH 7, black dashed; 4HFI, pH 4, blue dashed) are plotted. Mean distances for highest peaks have been labeled. Note: Distance distribution x-axes do not show non-adjacent distances that are beyond the sensitivity of our DEER experiments.

The distances between spin-labels on adjacent subunits attached at G60 (β2-β3 loop), T17 and N19 (β1), S46 and K48 (β2), S93C (β5), T99C and Q101C (β6), and I140C (β8) were shorter at pH 3.0 (desensitized state) compared to distances at pH 7.6 (**Fig. 5, Fig. S3**). The fact that we observed little to no proton-induced changes in the CW spectra at most positions (**Fig. 3**) while observing large distance changes with DEER spectroscopy suggests that there is a whole domain movement of this β-sheet upon proton-induced channel gating transitions, which does not alter the local conformations of the individual side chains. We performed error analysis on the background corrected and analyzed DEER distance distributions to validate the observed distance distributions (**Fig. S5**). The errors in the DEER distance distribution fits were small, indicating that the observed proton-induced distance changes were significant. The DEER derived distance distributions for D91R1 have more peaks than distributions with other sites, in part because the non-adjacent inter-subunit distances fall within the range of detection (**Fig. S3B**). The large number of peaks together with the shortening of the inter-subunit distance into the detectable range makes it difficult to discern and interpret an overall change in mean distance. However, the distribution clearly shifts towards shorter values when GLIC is in the desensitized state.

**Figure 5.**
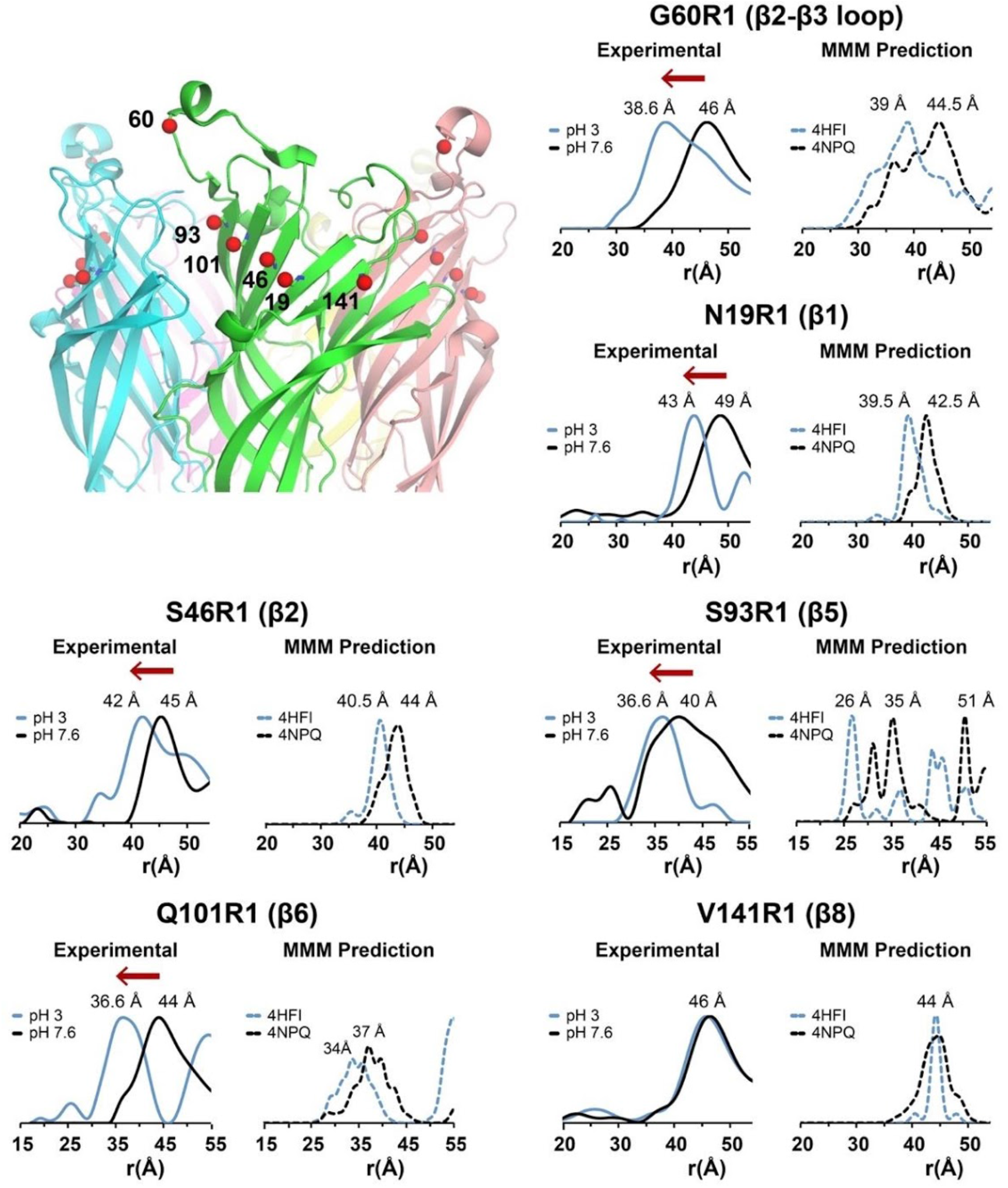
DEER derived distance changes for spin probes located on G60C (β2-β3 loop), N19C (β1), S46C (β2), S93C (β5), Q101C (β6) and V141C (β8). Top left panel: Expanded side view of GLIC ECD (PDB 4HFI) with C_β_ positions of spin-labeled residues depicted as red spheres. Experimental DEER interspin distance distributions showing the probability of a distance (P(r)) versus distance (r) are plotted at pH 7.6 (closed state, black) & pH 3.0 (desensitized state, blue). MMM simulated interspin distance distributions of corresponding residues showing the probability of a distance (P(r)) versus distance (r) obtained using crystal structures of GLIC (4NPQ, pH 7, black dashed; 4HFI, pH 4, blue dashed) are plotted. Mean distances for highest peaks have been labeled. Note: X-axes for distance distributions do not show non-adjacent distances. Spin labels attached to G60, N19, S46, S93 and Q101 move closer at pH 3.0. See **Fig. S3** for associated Q-band DEER dipolar evolution data.

We then compared our measured DEER distances from functional purified GLIC protein reconstituted in PE:PG liposomes with those calculated from the GLIC closed (4NPQ, pH 7) and open state crystal structures (4HFI, pH 4) after modeling MTSL side-chains at each position using Multiscale Modeling of Macromolecules (MMM (Polyhach, Bordignon et al. 2011); **Fig. 4**, **5**, **Fig. S3**). The *in-silico* distances from the closed and open state crystal structures predicted that, at most positions, the spin-probes move closer together in the open state, consistent with our experimental DEER data. V141R1 (β8) had no proton-induced change in DEER derived distances and the *in-silico* distance distributions also showed no change between the closed and open state structures (**Fig. 5**). Interestingly, the DEER derived experimental interspin distances for GLIC in the resting-closed state and desensitized state in liposomes were longer than the *in-silico* distances obtained from the closed and open state crystal structures, respectively (**Fig. 4F**, **Fig. S3C**). While some of the differences between experimental and *in-silico* computed distances may result from the rotameric MTSL library that we are using for our modeling, the data suggest that the structure of GLIC embedded in lipids is less compact than that predicted from crystal structures.

### Elastic Network Modeling

We used Elastic Network Modeling (ENM) to visualize the global structural motions in GLIC that would lead to the proton-induced decreases in the spin-label distances that we measured (**Fig. 6**). Since our DEER interspin distances at pH 7.6 were longer than those predicted from the presumed closed state GLIC crystal structure (4NPQ), we constructed a new resting state model for the GLIC ECD using our DEER-derived pH 7.6 distances and the closed state GLIC crystal structure (4NPQ) (GLIC-Mod7.6). We then constructed a model for the GLIC ECD in the desensitized state using GLIC- Mod7.6 and our DEER-derived distances at pH 3.0 (GLIC-Mod3.0). Comparing GLIC-Mod7.6 and GLIC-Mod3.0 showed that a global inward motion of the GLIC ECD accurately predicted the proton-induced decreases in DEER distances (**Fig. 6, Movie S1**).

**Figure 6.**
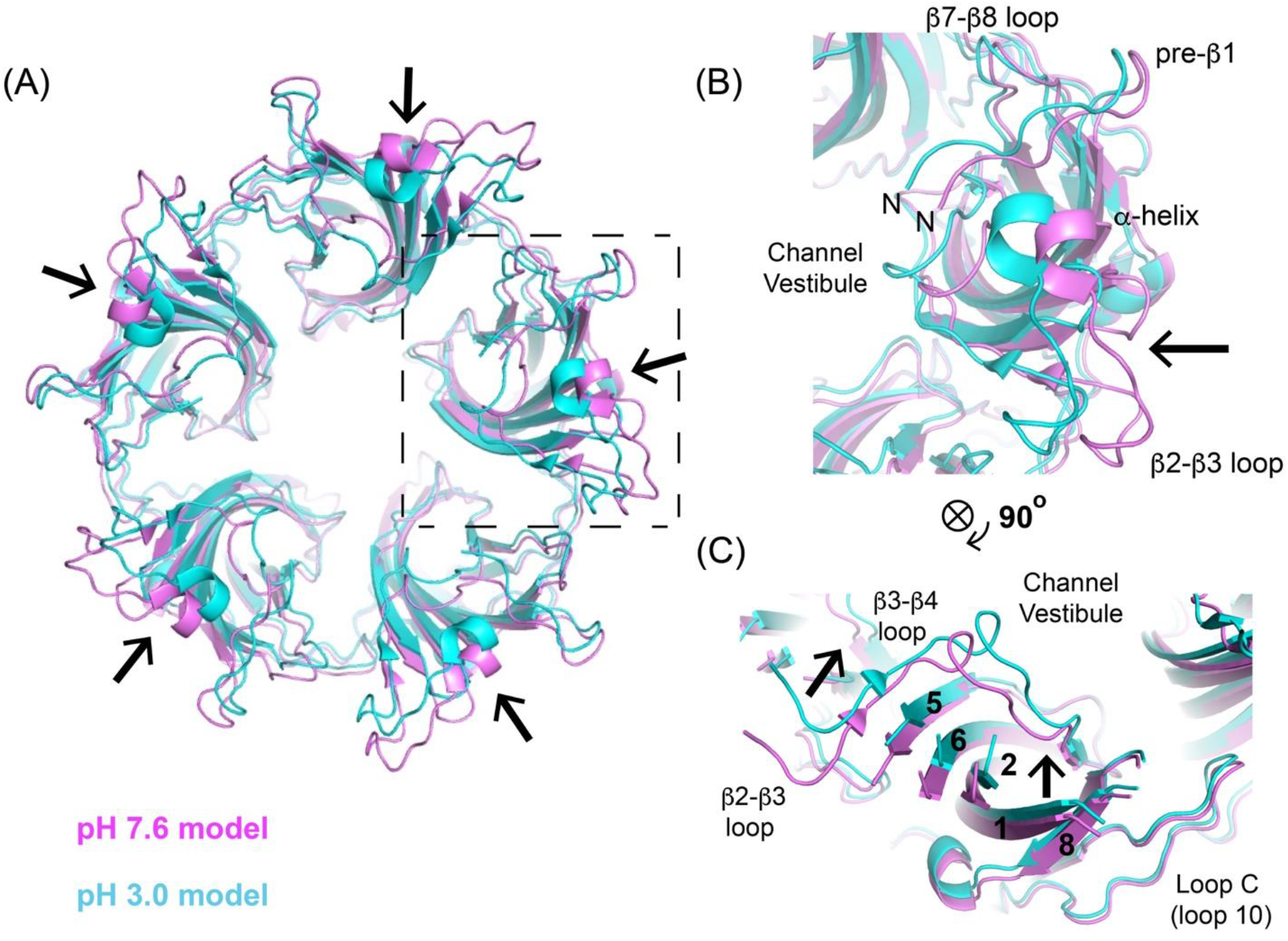
Inward tilting of ECD induced by proton binding. (A) Elastic network models of GLIC at pH 7.6 (magenta) and pH 3.0 (cyan). For clarity, only the ECD is shown. GLIC model at pH 7.6 (GLIC-Mod7.6) was generated using GLIC crystal structure at pH 7 (4NPQ) and experiment distance constraints from DEER at pH 7.6. GLIC model at pH 3.0 (GLIC-Mod3) was obtained using GLIC-Mod7.6 and experimental constraints from DEER at pH 3.0. Structures were aligned, and black arrows indicate the inward motion of GLIC inner β-sheet induced by proton binding. (B) Expanded view of one subunit highlighting inward motion of α-helix and β2-β3, β7-β8 loops and N-terminal pre-β1 region. (C) View rotated 90°clockwise from (B) highlighting motions of β5, β6, β2, β1 and β8 (black arrows). Parts of extracellular loops between β5-β6, β2-β3, β7-β8 and the pre-β1 N-terminal region have been hidden for clarity. See movie in supplementary materials for additional details (**Movie S1**).

### DEER spectroscopy reveals structural changes in the ECD of a gating-deficient GLIC mutant

Our EPR data indicates that the GLIC ECD adopts an inward tilted conformation when the protein is in a desensitized state in lipids. However, the data cannot distinguish when the tilt occurs. The ECD tilt could occur during a closed to open channel state transition and the ECD remains tilted in the desensitized state. Alternatively, the tilt could be associated specifically with a transition into a desensitized state from an open state. On the other hand, the tilt could occur during a closed to pre-active intermediate closed state transition. To help distinguish between these possibilities, we measured the distances between spin-labels attached at N19 (β1) in the GLIC ECD at pH 7.6 and pH 3.0 in the presence of a non-activating mutation Y251A (Y27’A) located in the M2-M3 loop. Previous studies have shown that Y251A (Y27’A) inhibits channel opening by stabilizing the protein in a ligand bound intermediate closed state (Gonzalez-Gutierrez, Cuello et al. 2013, Bertozzi, Zimmermann et al. 2016). *Xenopus* oocytes injected with GLIC N19C Y251A cRNA or purified GLIC N19R1 Y251A proteoliposomes displayed no proton-elicited currents, consistent with the Y251A mutation inhibiting GLIC double mutant channel opening. Interspin DEER derived distances at pH 7.6 for GLIC N19R1 in the Y251A background had a broad peak centered at 49 Å (**Fig. 7C)** similar to the distance measured in the absence of Y251A (**Fig. 5**). At pH 3.0, the interspin distance distributions decreased by ~5 Å (**Fig. 7C**), indicating that the spin probes attached to N19C moved closer at pH 3.0 even in the presence of a gating-deficient mutation. These data suggest that the proton-induced inward tilting of the GLIC ECD occurs during a resting, closed to pre-active intermediate closed state transition.

**Figure 7.**
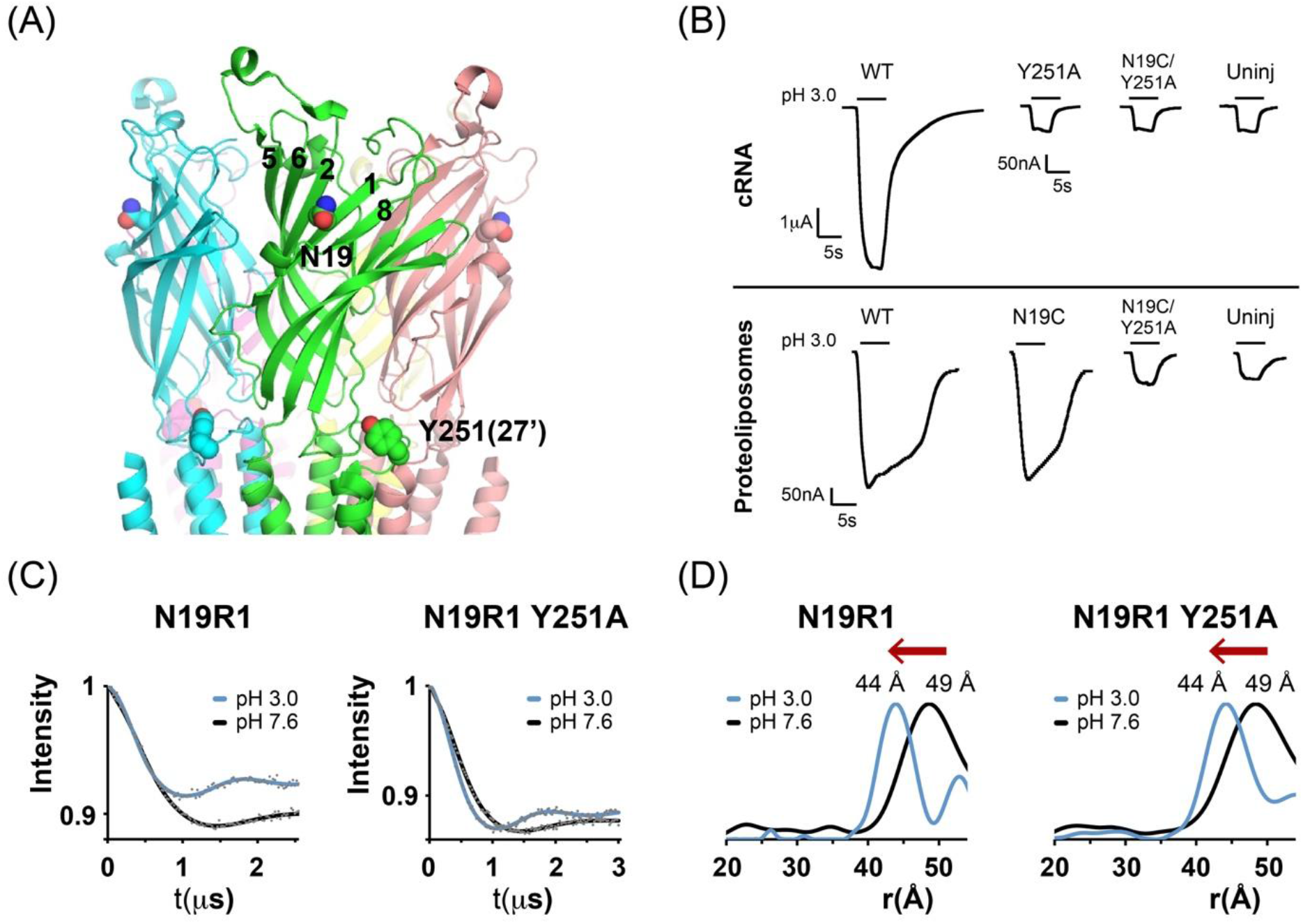
DEER derived distance changes for GLIC N19R1 in the presence of Y251A (Y27’A) mutation. (A) Side view of GLIC (PDB 4HFI) with N19 on β6 and Y251 (27’) on M2-M3 loop in space fill. (B) Macroscopic current responses to pH 3.0 applications of *Xenopus* oocytes injected with GLIC RNA and purified spin-labeled proteoliposomes. Currents from oocytes injected with GLIC N19C Y251A RNA and proteoliposomes were significantly smaller than those injected with wild type GLIC RNA and proteoliposomes (n =3) and not significantly different to those elicited from un-injected oocytes. (C) Comparison of background subtracted Q-band DEER dipolar evolution data (gray dots) of N19R1 and N19R1 Y251A fit using model-free fitting algorithms (pH 7.6, closed state, black solid lines; pH 3.0, desensitized state, blue solid lines) with the program LongDistances. (D) Corresponding experimental interspin distance distributions showing the probability of a distance (P(r)) versus distance (r) are plotted at pH 7.6 (black) & pH 3.0 (blue). Mean distances for highest peaks have been labeled.

## DISCUSSION

The conformational changes associated with pLGIC ligand-dependent gating transitions between closed, open and desensitized states remain poorly understood. Using SDSL EPR spectroscopy and functional GLIC channels reconstituted in liposomes, we studied changes in structure associated with gating transitions from a resting-closed state to a desensitized state. We targeted residues located at the top of β-strands 5, 6, 2, 1 and 8, which together comprise the inner β-sheet, to study proton-induced conformational changes in the ECD of GLIC. Our results reveal a significant structural rearrangement of the ECD inner β-sheet when GLIC transitions from the closed, resting state to the desensitized state. A proton-induced decrease in DEER derived mean inter-subunit distances occurred for residues located on all five β-strands: S93R1 (β5), T99R1 and Q101R1 (β6), K48R1 and S46R1 (β2), T17R1 and N19R1 (β1) and I140R1 (β8) and on the β2-β3 loop: G60R1 (**Fig. 4E, 5**, **Fig. S3B**). This structural change was also accompanied by a reorganization of the tertiary contacts of residues S93R1, D91R1 (β5), Q101R1 (β6) and I140R1 (β8) as revealed by their CW spectra (**Fig. 3**, **Fig. S2**). S93R1, D91R1 and Q101R1 located on the distal ends of β5 and β6 are more densely packed in the desensitized state (pH 3.0) than in the resting-closed state (pH 7.6). These residues are located at the (−) side of inter-subunit interface in proximity to loop C of the (+) subunit. Tight packing of this inter-subunit region suggests an increase in interaction between loop C of the (+) subunit and the distal ends of β5 and β6 of the (−) subunit during proton-mediated gating transitions. Indeed, previous EPR experiments on lipid reconstituted GLIC have reported an immobilization of loop C residues facing the β5 and β6 strands of the neighboring subunit in the desensitized state, supporting our conclusion (Velisetty and Chakrapani 2012). Overall, our EPR data demonstrate that proton-induced transition of GLIC from a resting-closed to a desensitized state causes significant motion of the ECD inner β-sheet, leading to a decrease in inter-subunit distances between residues located on β-sheets 5, 6, 2 and 1 as well as the β2-β3 loop, and this structural change likely involves a concerted displacement of these β-strands (**Fig. 3**).

We used ENM and our DEER derived distance data to predict the structural motions in the GLIC ECD that would fit our proton-induced distance changes. This analysis indicated that in lipids, the GLIC ECD in the desensitized state adopts a conformation where the inner β-sheet has moved inwards towards the central channel axis relative to its conformation in the resting-closed state (**Fig. 6, Movie S1**). The inward motion is more pronounced at β-strands 5, 6, 2, 1 and the β2-β3 loop, whereas the inward movement at β8 is relatively small. This structural change is similar to the inward tilting or “un-blooming” motion of the ECD predicted by comparing the closed and open crystal structures of GLIC and GluCl (Sauguet, Shahsavar et al. 2014, Nemecz, Prevost et al. 2016). Comparing closed and open state structures of GLIC, GluCl and the α1-glycine receptor also predict a quaternary counter-clockwise twisting motion of the ECD during channel opening. However, a global rotation will not change the intersubunit distances and therefore would not be detected in our experiments.

Comparison of the closed and open structures of GluCl and α1-glycine receptor also predicts that channel opening is associated with an outward movement of the ends of β7, β8 and β10 near the ECD-TMD interface, while comparison of GLIC structures predicts that this region moves slightly inwards when the channel opens. To explore conformational changes in this region, residues S123 (β7) and S191 (β10) located near the ECD-TMD interface of GLIC were mutated to cysteine and probed for proton-induced structural changes using CW EPR and DEER spectroscopy (**Fig. S4A**). DEER derived interspin distances at pH 7.6 and pH 3.0 show that S123R1 and S191R1 moved closer at pH 3.0, suggesting that the ends of the outer β-sheet near the ECD-TMD interface move inwards on proton induced gating transitions from a closed state to a desensitized state (**Fig. S4B, C**). This suggests that the hinge for the inward tilt of GLIC ECD is near the ECD-TMD interface.

To examine when the ECD inward tilting motion occurred in the gating cycle of GLIC, we probed for proton-induced ECD conformation changes in a non-activating GLIC mutant: GLIC Y251A (Y27’A). The Y251A mutation in the M2-M3 loop impairs gating but not ligand binding, and has thus been suggested to stabilize the protein in a liganded-closed channel intermediate state (Gonzalez-Gutierrez, Cuello et al. 2013, Bertozzi, Zimmermann et al. 2016). The proton-induced decrease in mean DEER derived distances detected at N19R1 (β1) in the presence of Y251A was identical to that measured for GLIC N19R1, suggesting that the GLIC ECD has adopted an inward-tilted conformation when the channel is in a liganded-closed pre-active state (**Fig. 7**). Data obtained from a variety of pLGICs support the idea that GLIC Y251A (Y27’A) stabilizes a ligand-bound closed state and not a desensitized state. Kinetic analyses of single channel currents from nAChR and GlyRs show that mutations in M2-M3 loop reduce the liganded gating equilibrium constant without affecting transitions to a desensitized state (Lewis, Sivilotti et al. 1998, Grosman, Salamone et al. 2000). The protein conformation observed in the crystal structure of GLIC Y251A is identical to the “locally closed” GLIC crystal structure (Gonzalez-Gutierrez, Cuello et al. 2013). This “locally closed” conformation of GLIC has been suggested to represent a closed channel intermediate in the transition of GLIC from the closed to open state (Prevost, Sauguet et al. 2012, Sauguet, Shahsavar et al. 2014).

In conclusion, our EPR data suggest that an inward tilting of GLIC ECD is associated with the transition of the channel from a resting-closed state to a ligand-bound closed channel intermediate state, and that the ECD remains in this conformation when the channel transitions to the open and the desensitized states (**Fig. 8**). The inward tilting motion was initially reported by molecular simulations of GluCl relaxation from an open (ivermectin-bound) state to a closed channel state on the removal of ivermectin, and it was proposed that this structural rearrangement is crucial in coupling changes at the agonist-binding site in the ECD to the TMD pore (Calimet, Simoes et al. 2013, Taly, Hénin et al. 2014). Single-channel electrophysiology experiments on nAChRs and GlyRs also suggest that binding of agonist stabilizes the channel in a pre-activated intermediate “flip” state from which the channel eventually transitions into an open state (Lape, Colquhoun et al. 2008, Nemecz, Prevost et al. 2016). Thus, the inward-tilting motion of the ECD observed in our experiments could represent an early structural change in the gating cycle that is associated with transition of the receptor to a high-affinity binding conformation. Recent fluorescence-quenching experiments on bimane-labeled GLIC also reported a global compaction of the ECD during proton-induced gating transitions and suggested that this motion is associated with a pre-active intermediate state (Menny, Lefebvre et al. 2017). String method molecular simulations of GLIC deprotonation have also identified a closed-channel intermediate state with the ECD stabilized in a compact conformation (Lev, Murail et al. 2017).

**Figure 8.**
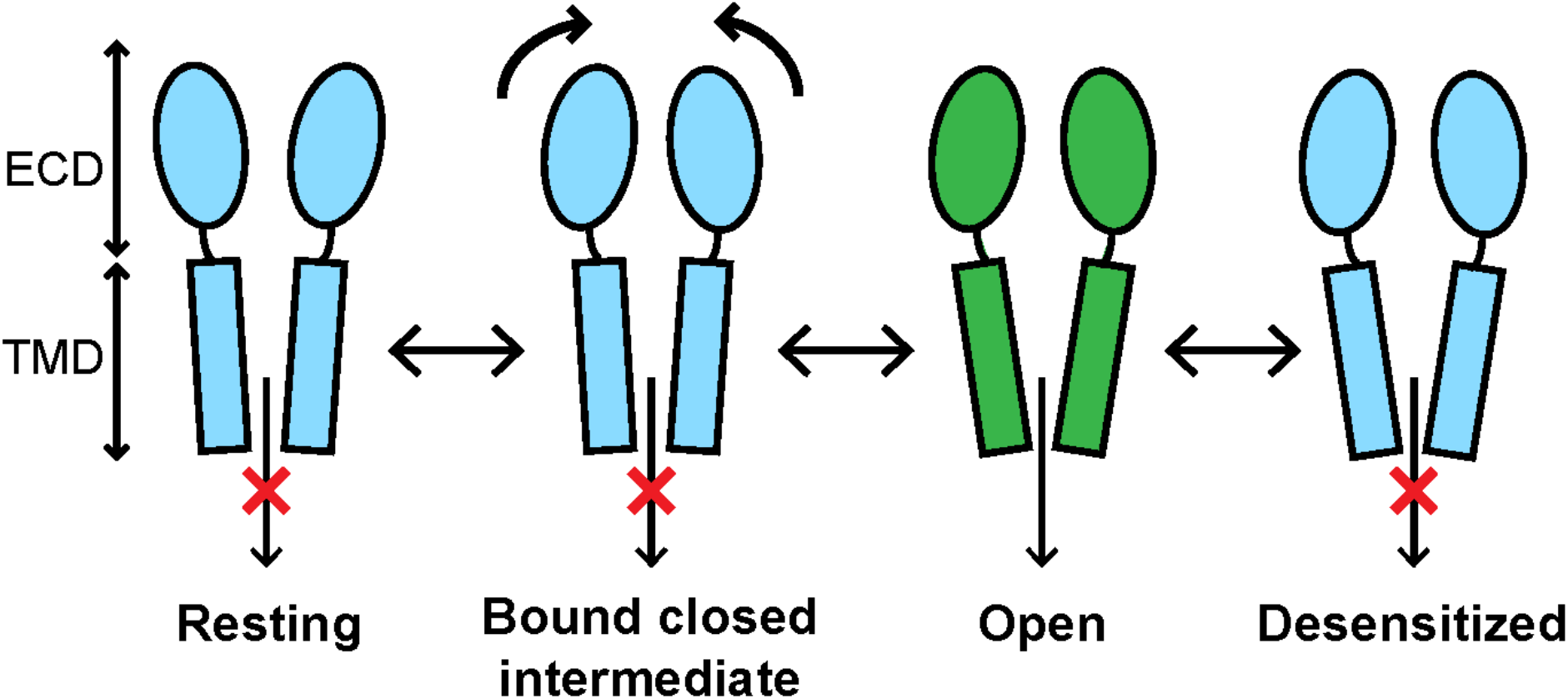
Schematic representation of ECD inward tilting motion during GLIC gating cycle. The inward tilting motion of GLIC ECD is associated with channel transition from a resting closed state to a ligand bound pre-activated closed state.

Our data also indicate that there is no significant change in the position of the ECD during the open to desensitized state transition. Voltage-clamp fluorometry experiments on α1-GlyR and GABA_A_Rs have also reported that residues near the agonist-binding site in the ECD do not show any additional conformational changes during desensitization (Muroi, Czajkowski et al. 2006, Akk, Li et al. 2011, Wang and Lynch 2011, Keramidas and Lynch 2013). Our data thus support a desensitization mechanism where the ECD remains in an inward-tilted high-affinity liganded conformation and channel closure is achieved by structural changes in the TMD (Morales-Perez, Noviello et al. 2016, Nemecz, Prevost et al. 2016, Gielen and Corringer 2018, Laverty, Desai et al. 2019, Masiulis, Desai et al. 2019). Further experiments are needed to fully understand the protein motions that stabilize the receptor in a high-affinity desensitized state.

## MATERIALS AND METHODS

### Cloning and mutagenesis

The GLIC DNA gift from Dr. Raimund Dutzler (University of Zurich) was subcloned into pET-26b vector (Novagen) for bacterial expression. This construct has an N-terminal *pelB* signal sequence followed by a (His)_10_ –tag, DNA sequence for maltose-binding protein (MBP), and the GLIC DNA sequence (residues 44-359). We modified the original ~20 amino acid linker sequence between MBP and GLIC to introduce a Tobacco Etch Virus protease (TEV protease) cleavage site (ENLYFQG) at the N-terminus of the GLIC sequence. Mutations were created using the QuikChange site-directed mutagenesis method with *PfuUltra* II Fusion HS DNA polymerase (Agilent) and a single mutagenesis primer. Reaction product was digested with *Dpn*I (FastDigest, ThermoFisher) and transformed into in-house *E. coli* Top10F’ cells. Positive clones were selected, and the presence of the mutation was confirmed by DNA sequencing.

### Protein expression and purification

*E. coli* OverExpress C43 (DE3) cells (Lucigen) cells were transformed with the pET-26b vector containing the DNA of GLIC mutants. Cells were grown in LB medium containing 50 μg/mL Kanamycin at 37°C to an OD_600_~1.0 (as measured in a Beckman Coulter DU 640 spectrophotometer) and then quickly transferred on ice to cool for ~20 mins. Protein expression was induced by adding 0.1 mM isopropyl β-D-1-thiogalactopyranoside (IPTG) and incubating the cells for ~16 hr at 20°C. All following steps were performed at 4°C unless noted otherwise. Cells were harvested by centrifugation and lysed using EmulsiFlex-C5 (Avestin) after resuspension in 20 mM Tris, 150 mM NaCl, pH 7.6 (Buffer B1) supplemented with 2 μM pepstatin, 2 μg/μL leupeptin, 1 mM ethylenediaminetetraacetic acid (EDTA), 1mM phenylmethanesulfonyl fluoride (PMSF). Cell lysate was cleared by centrifugation at 4000 rpm in JA14 rotor (Beckman Coulter) for 40 min and the supernatant was collected and spun at 35,000 rpm in 50.2Ti rotor (Beckman Coulter) for 1 hr to pellet membranes. Membranes were resuspended in Buffer B1 supplemented with 15 mM imidazole, 5 μg/μL leupeptin and 2% n-dodecyl-β-D-maltopyranoside (DDM, Anatrace), mixed using a Dounce homogenizer and incubated for ~16 hr at 4°C with agitation. DDM solubilized membranes were cleared by centrifugation at 45,000 rpm in 50.2Ti rotor (Beckman Coulter) for 45 min and the supernatant was purified using affinity chromatography with a Ni-NTA column (Qiagen). Ni-NTA column bound with MBP-GLIC fusion protein was washed with 5 column volumes of Buffer B1 with 50 mM imidazole and 1% DDM, followed by 5 column volumes of Buffer B1 with 50 mM imidazole and 0.02% DDM. MBP-GLIC was eluted using 2 column volumes of Buffer B1 with 300 mM imidazole and 0.02% DDM and then applied to an amylose resin column (New England Biolabs). Amylose resin column bound to MBP-GLIC was washed with 2 column volumes of Buffer B1 with 0.02% DDM and MBP-GLIC was eluted using 2 column volumes of Buffer B1 with 20mM maltose and 0.02% DDM. TEV protease (in-house purified) was added to the purified MBP-GLIC fusion protein in the ratio TEV: MBP-GLIC = 1: 20 (by weight) and the sample was agitated for 12-14 hr at 4°C to cleave the (His)_10_-MBP protein. The TEV digested sample was applied to a Ni-NTA column (Qiagen) to remove the (His)_10_-MBP and (His)_6_-containing TEV protease. Purified GLIC was collected in the column flow through and by additional elution using 2 column volumes of Buffer B1 with 30 mM imidazole and 0.02% DDM. Purified GLIC pentamer was first treated with 25-fold molar excess of DTT for 5 min at room temperature, followed by addition of 100-fold molar excess of 2,2,5,5-tetramethylpyrroline-3-yl-methanethiosulfonate spin label (MTSL, Toronto Research Chemicals) to specifically label the cysteines. MTSL-labeled GLIC was concentrated using Amicon Ultra-15 (100 kDa molecular weight cut off) concentrator tubes (EMD Millipore) and centrifuged at 90,000 rpm in TLA-120.2 rotor (Beckman Coulter) for 1 hr to remove protein aggregates. The protein in the supernatant was collected and subjected to gel filtration chromatography in a Superose6 10/300 GL column (GE Healthcare) previously equilibrated in Buffer B1 with 0.02% DDM. Peak fractions corresponding to pentameric GLIC (~182 kDa) were collected, pooled and concentrated to 2-4 mg/mL using Amicon Ultra-4 (100 kDa molecular weight cut off) concentrator tubes (EMD Millipore), flash frozen in liquid nitrogen and stored at −80°C.

### Reconstitution of purified GLIC into liposomes

Purified GLIC protein was reconstituted into liposomes formed with 1-palmitoyl-2-oleoyl-sn-glycero-3-phosphoethanolamine (PE, Avanti Polar Lipids) and 1-palmitoyl-2-oleoyl-sn-glycero-3-phospho-(1’-rac-glycerol) (PG, Avanti Polar Lipids) at a PE: PG = 2.7:1 molar ratio. Lipid mixtures at a concentration of 20 mg/mL in chloroform were mixed in appropriate volume, dried with argon and kept under vacuum overnight. Dried lipids were hydrated with appropriate volume of Buffer B1 and sonicated to form liposomes at a final concentration of 40 mM. PE: PG liposomes were mixed with GLIC purified in DDM, typically at 4,000-fold molar excess. The protein: lipid mixture was incubated for 3 hr at 4°C, following which the sample was diluted 2-fold in Buffer B1 containing 10% glycerol and incubated overnight at 4°C. To remove DDM, around 200 mg of prewashed Biobeads (BioRad) were added to the sample for 8-9 hr under gentle agitation and then removed. GLIC proteoliposomes were collected by centrifugation of the Biobead-free sample at 100,000 rpm in TLA-120.2 rotor (Beckman Coulter) for 1 hr, and the pellets were stored at −80°C.

### Two-Electrode Voltage Clamp Recording of *Xenopus* oocytes

#### cRNA injection

Capped cRNAs encoding WT and mutant GLIC were transcribed *in vitro* using the mMessage mMachine T7 kit (Ambion). Oocytes were obtained from an in-house *X. laevis* colony or from Ecocyte Biosciences. Single oocytes were injected with 27 nL of cRNA (100 ng/μL). Injected oocytes were incubated at 16°C in ND96 (5 mM HEPES pH 7.4, 96 mM NaCl, 2 mM KCl, 1 mM MgCl_2_, 1.8 mM CaCl_2_) supplemented with 100 μg/ml of gentamycin and 100 μg/mL of bovine serum albumin for 2–4 days before use for electrophysiological recordings. Oocytes were perfused continuously with ND96 at pH 7.4 at a flow rate of 5 mL/min in a 200 μL bath while being held under two-electrode voltage clamp at −40 mV. Borosilicate glass electrodes (Warner Instruments) used for recordings were filled with 3 M KCl and had resistances of 0.4 to 1.0 MΩ. Electrophysiological data were collected using Oocyte Clamp OC-725A (Warner Instruments) interfaced to a computer with a Digidata 1440A (Axon CNS) and were recorded using the AxoScope program, version 10.2 (Molecular Devices). Proton-induced currents were measured by perfusing ND96 buffered at pH 7.0-3.0. pH 4.6-3.0 buffers used 5mM Na Citrate to replace HEPES as the buffering agent. Proton-induced currents were measured at pH 3.0 until peak current amplitudes varied by <10%, then measured at pH 7.0 as a control, followed by a final application of pH 3.0. pH-induced currents from uninjected oocytes were used as controls.

#### Proteoliposome injection

Pellets of purified GLIC mutants reconstituted into liposomes were thawed on ice. The amount of protein in a pellet was estimated by assuming 70% reconstitution efficiency. The pellets were resuspended in buffer B1 to a protein concentration of approximately 1 mg/mL and were subjected to two rounds of freeze-thaw. To facilitate the injection of this somewhat viscous solution, oocytes were injected with a glass pipet with the diameter adjusted to around 5-10 μm. Protein-injected oocytes were incubated for 16-24 h at 16°C before recording. Two-electrode voltage clamp of oocytes injected with lipid reconstituted GLIC protein was performed in the same manner as oocytes injected with GLIC cRNA. Dose-response traces were generated using pH 6.4, 4.6, 3.5 and 3.0.

### EPR Spectroscopy

Continuous wave (CW) EPR spectroscopy was carried out at room temperature on a Bruker ELEXSYS 500 X-band spectrometer equipped with a super high Q (SHQ) cavity (Bruker Biospin). Upon change in pH to 3.0, the proteolipid samples were frozen in liquid nitrogen and thawed on ice to ensure even distribution of the protons inside and outside the vesicles. Spectra were then recorded over 100 G under non-saturating conditions with a 100 kHz field modulation of 1.5 G. The DEER spectroscopy experiments were carried out at the National Biomedical EPR Center using a Bruker ELEXSYS 580 Q-band spectrometer equipped with an EN5107D2 resonator (Bruker Biospin). Samples were typically 12 μL in volume at a pentamer concentration of ~30 μM, contained 20% deuterated glycerol as a cryoprotectant, were flash frozen using a dry ice-acetone slurry, and run at 80 K. A four-pulse DEER sequence was used to collect the dipolar evolution data (Martin, Pannier et al. 1998, Pannier, Veit et al. 2000). The resulting data were background corrected and analyzed for distance distributions using the LongDistances software written by C. Altenbach (UCLA) (Toledo Warshaviak, Khramtsov et al. 2013). Errors of the distance distribution fits were calculated from 100 iterations of the error analysis machinery implemented in the LongDistances software and plotted at ± 1 standard deviation (**Fig. 2.13**). The upper reliable distance limit (d) based on the maximum data collection time (t) recorded for these data sets (d ≈ 5(t/2)1/3; (Jeschke 2012) is represented by the x-axes of the distance distribution plots shown.

### Computational modeling of MTSL and experimental DEER distances on GLIC crystal structures

*In-silico* modeling of MTSL on GLIC was done using the rotamer library approach implemented in the software MMM version 2015.1 (Multiscale Modeling of Macromolecules (Polyhach, Bordignon et al. 2011), http://www.epr.ethz.ch/software.html). Crystal structures of GLIC in the presumed closed (pH 7, PDB 4NPQ) and open state (pH 4, PDB 4HFI) were used as input templates. Each target residue was selected in all five subunits and rotamer computation was performed at a temperature of 175 K (cryogenic) with MTSL as the spin-label. Distance distributions were calculated between MTSL rotamers attached to the target residue on all five GLIC subunits. Structural models for GLIC ECD in the resting-closed (GLIC_Mod7.6) and desensitized (GLIC_Mod3) states in liposomes were made using Elastic network modeling (ENM) implemented in MMM by fitting experimental DEER distance constraints to the GLIC crystal structures (Jeschke 2012). In detail, the GLIC crystal structure in the presumed closed state (4NPQ, pH 7) was selected as the starting template and fitted to experimental DEER distances at pH 7.6 to generate GLIC_Mod7.6. GLIC_Mod7.6 was then fitted to the experimental DEER distances at pH 3 to generate GLIC_Mod3. Mean values of the highest adjacent interspin distance peak for residues T17R1, N19R1, S46R1, K48R1, G60R1, S93R1, T99R1, Q101R1, I140R1, V141R1, S123R1 and S191R1 derived experimentally using DEER at pH 7.6 and pH 3.0 were used as distance constraints for ENM. The distance constraint files used for ENM are available in the Supplementary Information. The morph between GLIC_Mod7.6 and GLIC_Mod3 was generated using UCSF Chimera (Pettersen, Goddard et al. 2004) and all figures were prepared using Pymol (Schrodinger 2015).

---

**Figure S1.**
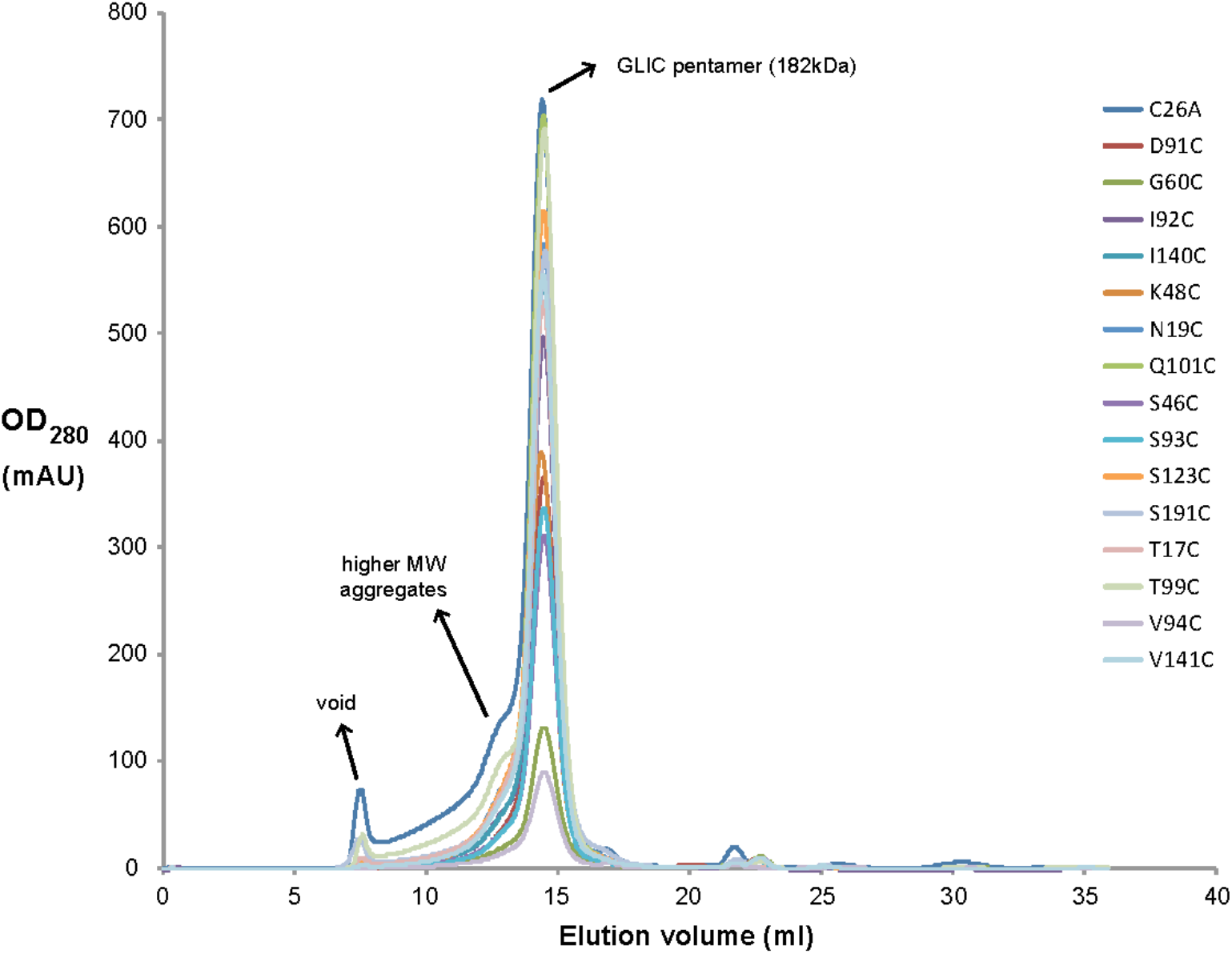
Size exclusion chromatography of purified GLIC protein. Peak elution profiles of DDM solubilized GLIC mutants run on Superose 6 10/300 GL column (GE Healthcare) show a monodisperse protein population corresponding to the GLIC pentamer (182 kDa). Only the peak fractions corresponding to the GLIC pentamer were collected and used for lipid reconstitution and EPR studies.

**Figure S2.**
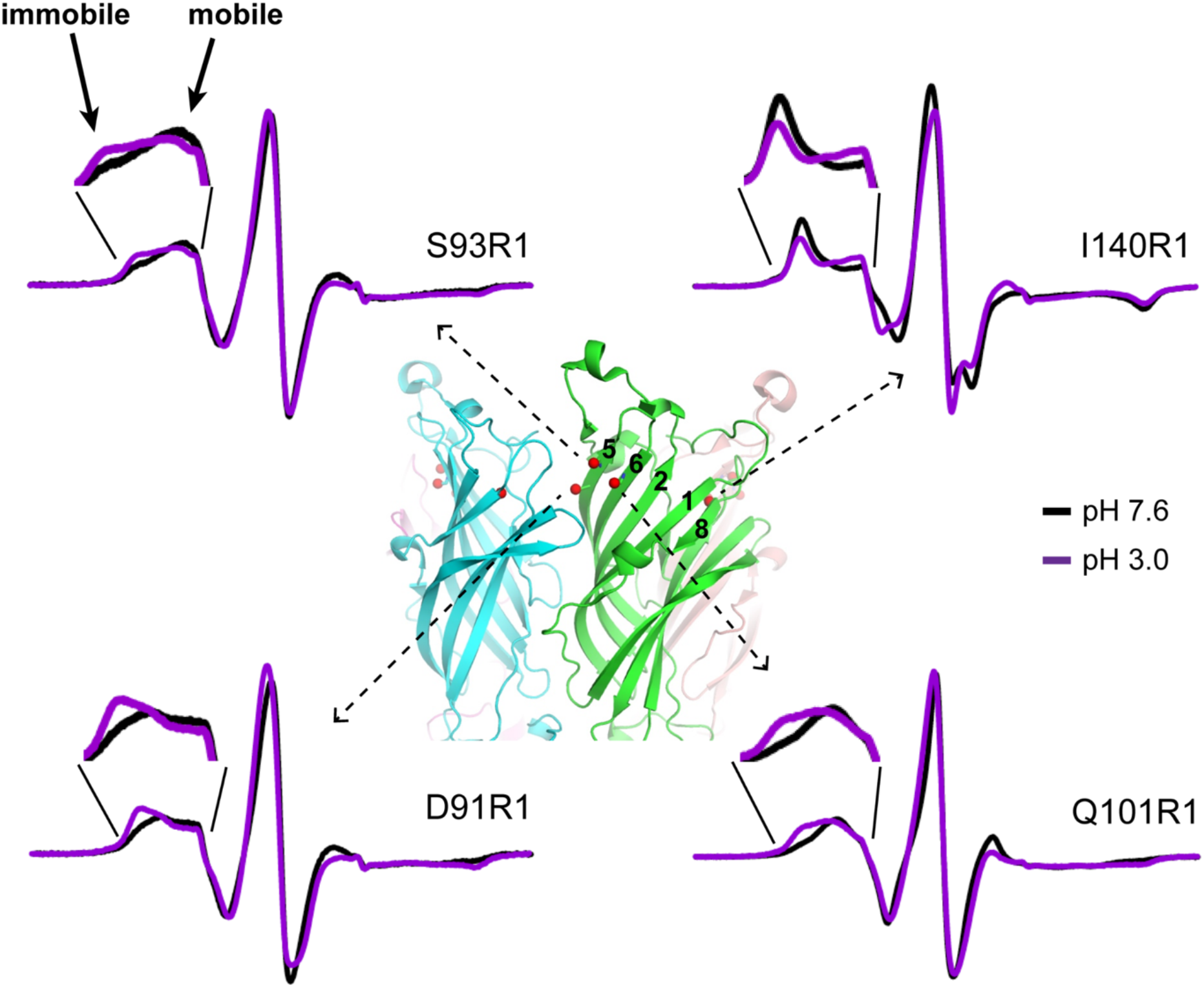
Proton-induced CW spectral changes for GLIC D91R1, S93R1, Q101R1 and I140R1. Comparison of X-band CW EPR spectra of spin-labeled GLIC protein at pH 7.6 (black, closed state) and pH 3.0 (magenta, desensitized state). The low-field regions of the spectra are enlarged to highlight proton-induced changes. Mobile and immobile components in the low-field region of S93R1 spectrum are indicated. Center: Side view of GLIC ECD (PDB 4HFI) with C_β_ atoms of D91, S93 (β5), Q101 (β6) and I140 (β8) displayed as red spheres and β-strands labeled. I140 on β8 faces the channel central symmetry axis.

**Figure S3.**
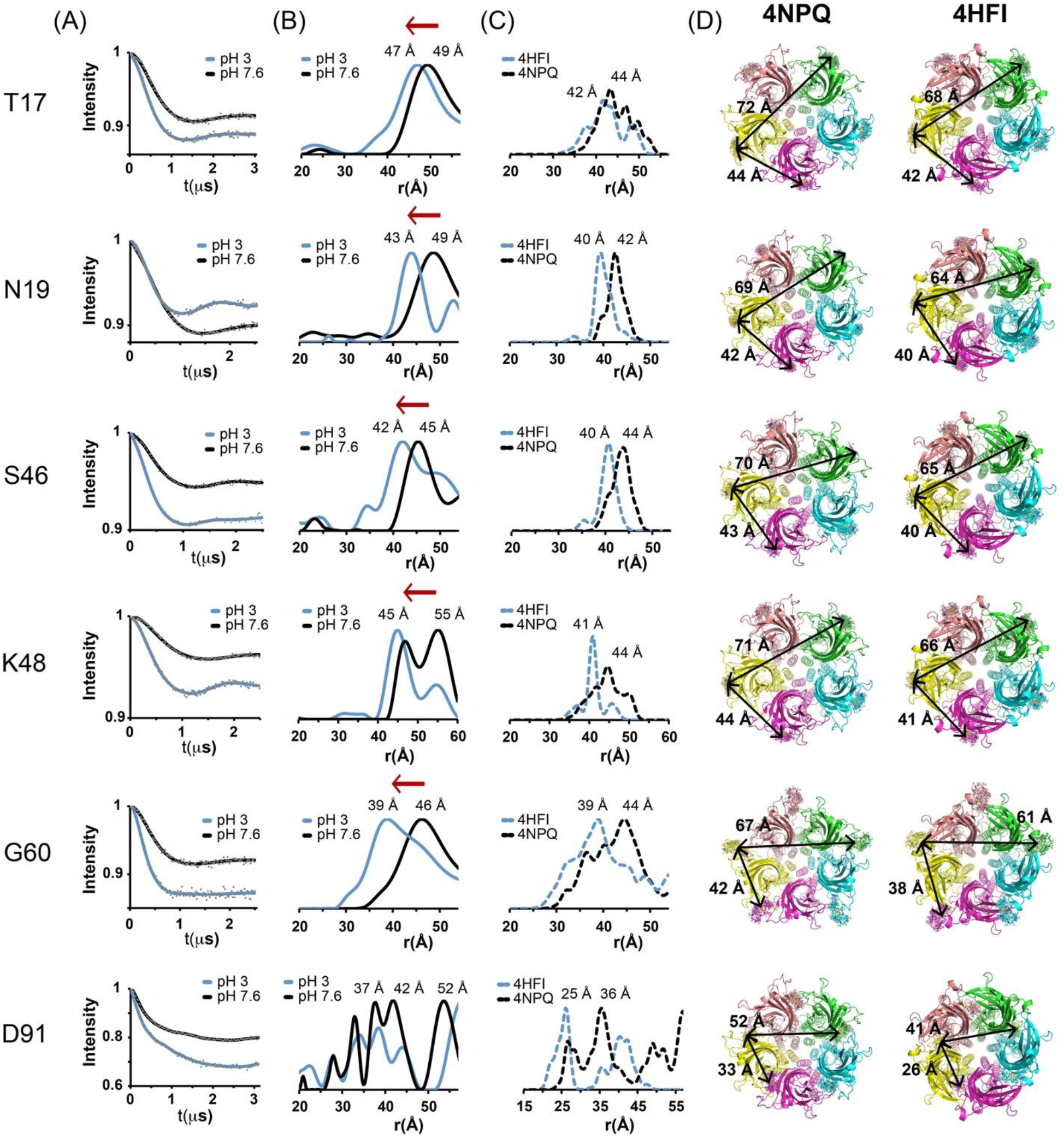
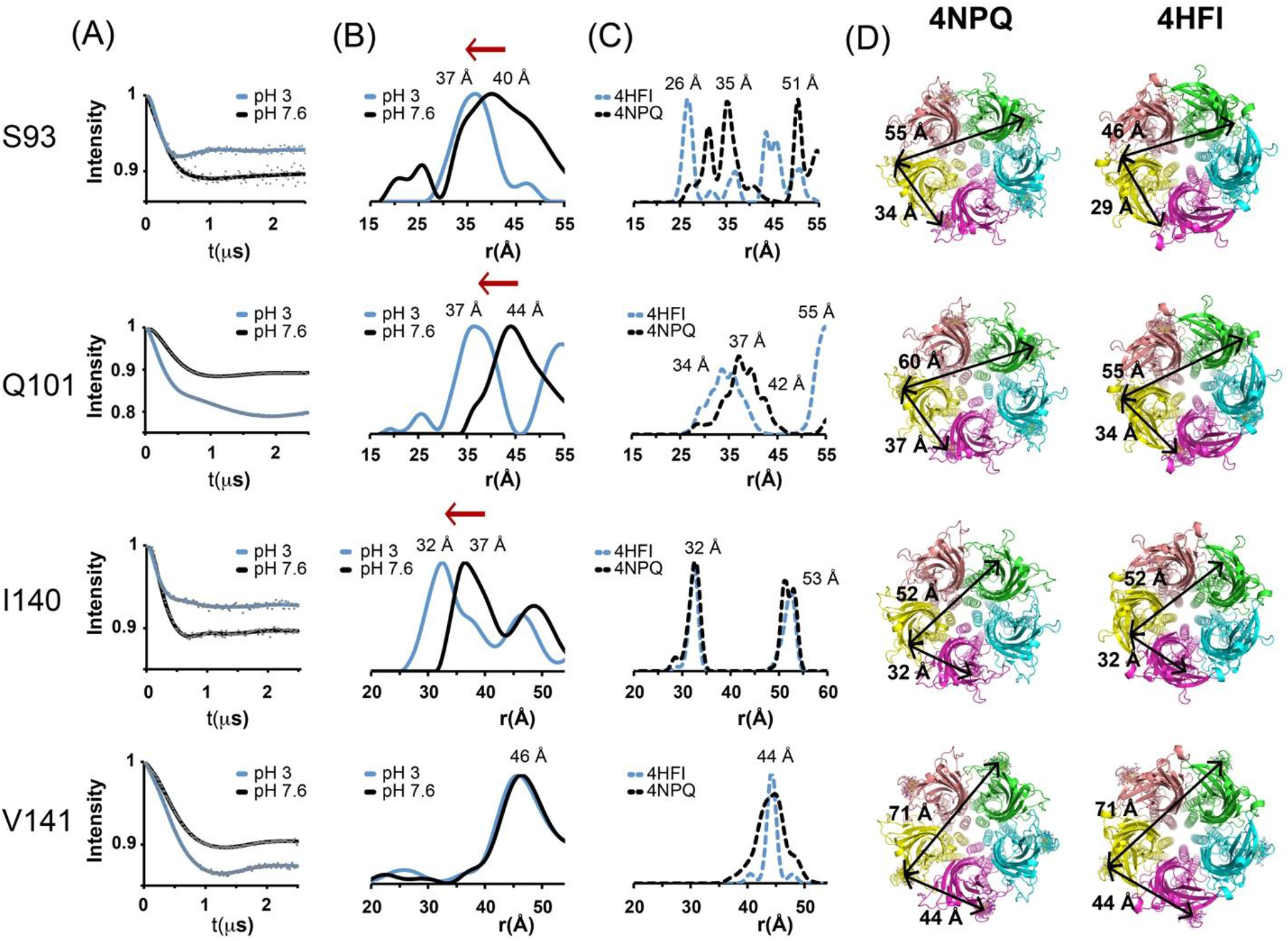
Summary of DEER spectroscopy data. (A) Background corrected Q-band DEER dipolar evolution data (gray dots) of spin-labeled GLIC protein fit using model-free fitting algorithms at pH 7.6 (closed state, black solid lines) and pH 3.0 (desensitized state, blue solid lines) with the program LongDistances. (B) Corresponding experimental interspin distance distributions showing the probability of a distance (P(r)) versus distance (r) are plotted at pH 7.6 (black) and pH 3.0 (blue). Mean distances for highest peaks are labeled. (C) MMM simulated interspin distance distributions for respective residues showing the probability of a distance (P(r)) versus distance (r) obtained from crystal structures of GLIC (4NPQ, pH 7, black dashed lines; 4HFI, pH 3, blue dashed lines). Mean distances for highest peaks are labeled. Note: Distance distribution x-axes do not show non-adjacent distances that are beyond the sensitivity of our DEER experiments. (D, E) Top-down views of GLIC (4NPQ, pH 7; 4HFI, pH 4) showing predicted spin label rotamers displayed in sticks. Black lines depict average interspin distances between adjacent and non-adjacent subunits obtained using MMM.

**Figure S4.**
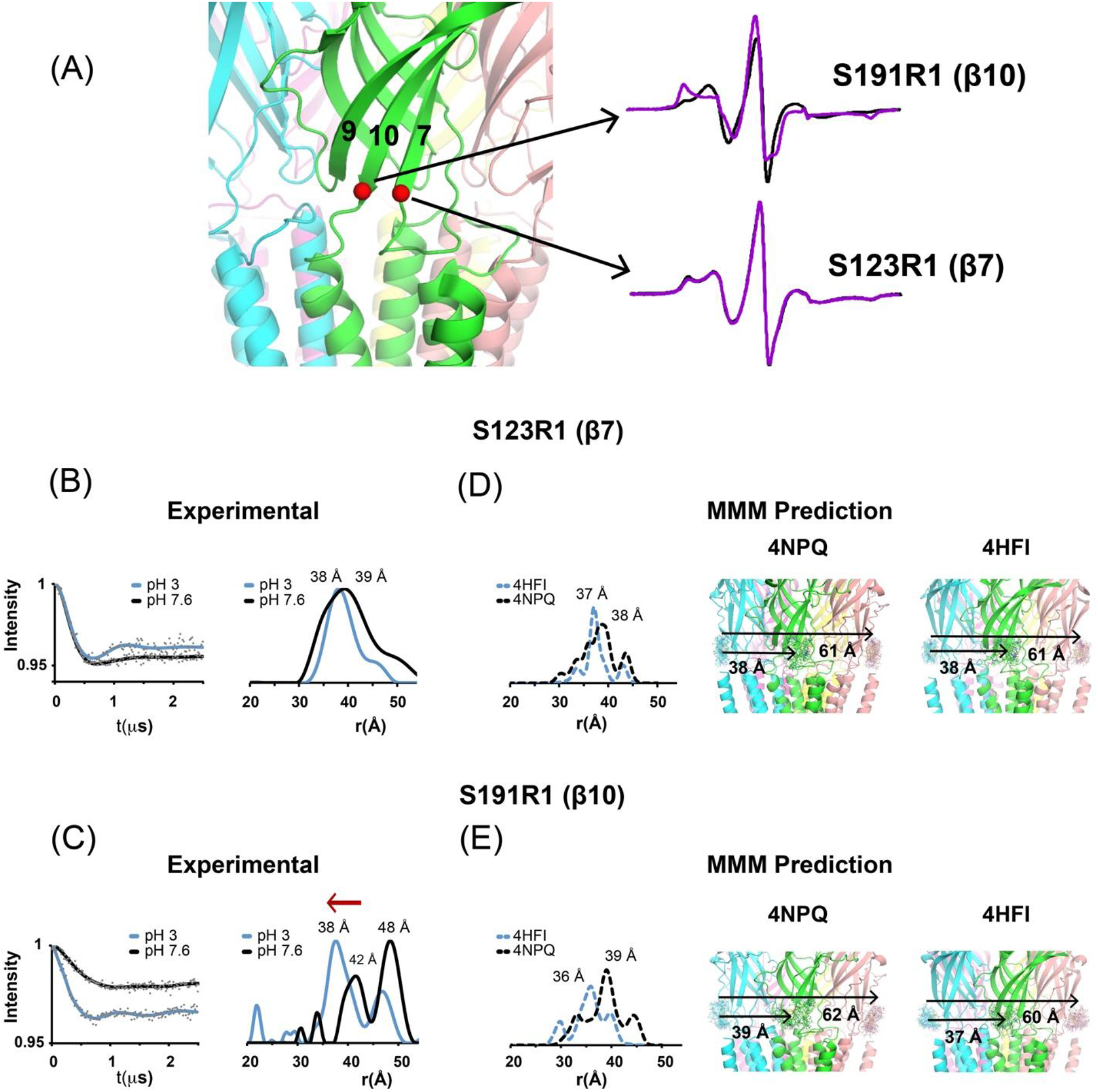
CW EPR and DEER spectroscopy data for residues S123R1 (β7) and S191R1 (β10). (A) Side view of GLIC (PDB 4HFI) with Cβ atoms of S123 on β7 and S191 on β10 displayed as red spheres. Comparison of CW EPR spectra of spin-labeled GLIC protein at pH 7.6 (black, closed state) and pH 3.0 (magenta, desensitized state). (B, C) Background corrected Q-band DEER dipolar evolution data (gray dots) of spin-labeled GLIC protein fit using model-free algorithms at pH 7.6 (closed state, black solid lines) and pH 3.0 (desensitized state, blue solid lines) with the program LongDistances. Corresponding experimental interspin distance distributions showing the probability of a distance (P(r)) versus distance (r) are plotted for pH 7.6 (black) and pH 3.0 (blue). Mean distances for highest peaks are labeled. (D, E) MMM simulated interspin distance distributions for respective residues showing the probability of a distance (P(r)) versus distance (r) obtained from crystal structures of GLIC (4NPQ, pH 7, black dashed lines; 4HFI, pH 3, blue dashed lines). Mean distances for highest peaks are labeled. Note: Distance distribution x-axes do not show non-adjacent distances that are beyond the sensitivity of our DEER experiments. Side views of GLIC (4NPQ, pH 7; 4HFI, pH 4) showing predicted spin label rotamers displayed in sticks. Black lines depict average interspin distances between adjacent and non-adjacent subunits obtained using MMM (two subunits in back are hidden for clarity).

**Figure S5.**
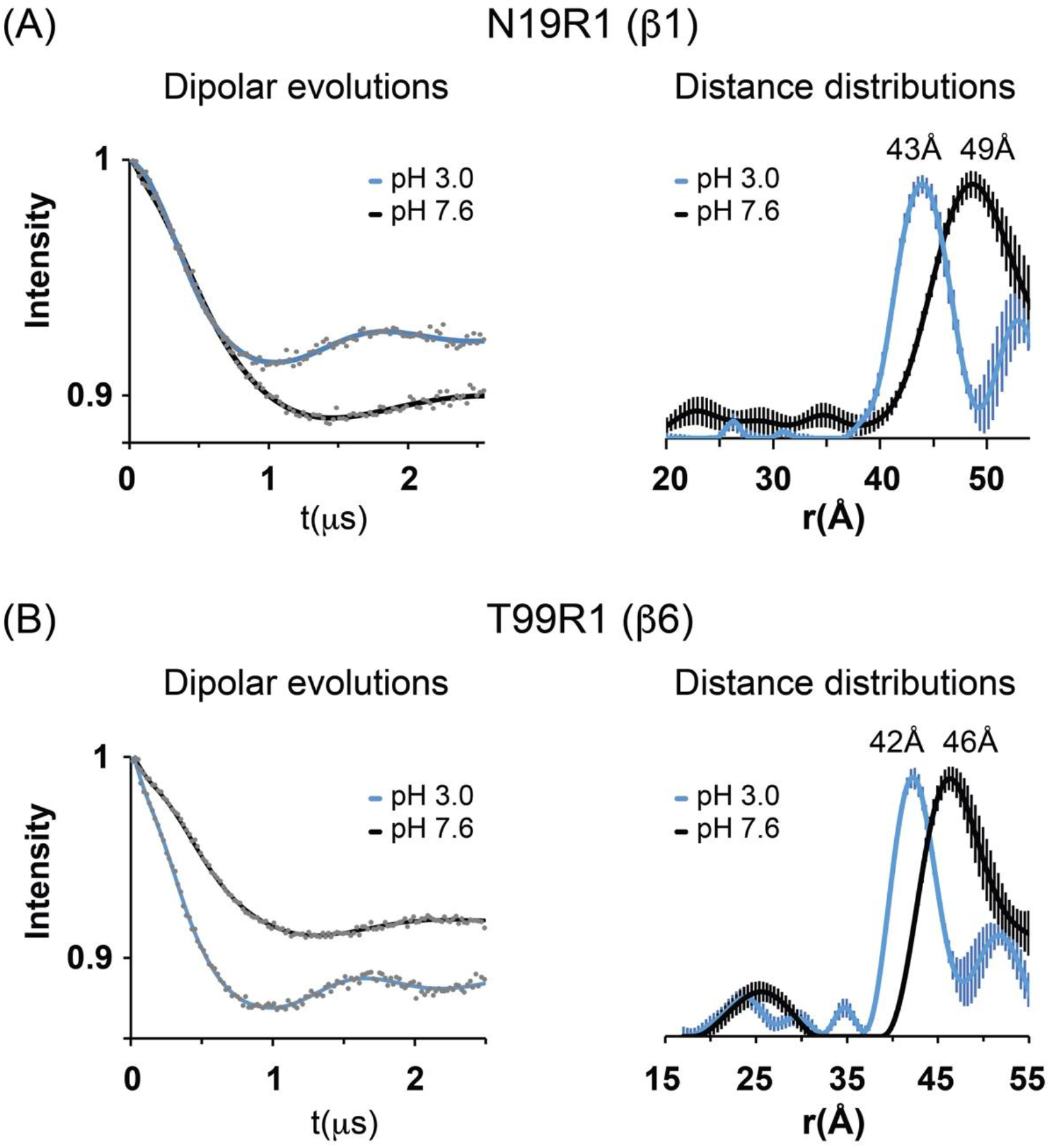
DEER spectroscopy data for residues N19R1 (β1) and T99R1 (β6). Background corrected Q-band DEER dipolar evolution data (gray dots) of spin-labeled GLIC mutants N19C (β1) and T99C (β6) fit using model-free algorithms at pH 7.6 (closed state, black solid lines) and pH 3.0 (desensitized state, blue solid lines) with the program LongDistances. Corresponding experimental interspin distance distributions showing the probability of a distance (P(r)) versus distance (r) are plotted for pH 7.6 (black solid line) and pH 3.0 (blue solid lines) with the respective errors (± 1 standard deviation) as computed using LongDistances program shown with vertical lines. Mean distances for the highest peaks are labeled. Note: Distance distribution x-axes do not show non-adjacent distances that are beyond the sensitivity of our DEER experiments.

## ACKNOWLEDGEMENTS

We are grateful to Dr. Raimund Dutzler (University of Zurich) for providing us with the GLIC DNA. We would also like to thank Dr. Vadim Klenchin for his help with GLIC purification.

## AUTHOR CONTRIBUTIONS

CC - Experimental design, data analysis, manuscript writing, manuscript editing, project management ; CK - EPR spectroscopy experiments, data analysis, manuscript editing; JB - two-electrode voltage clamp experiments, manuscript editing, data analysis, AS - mutagenesis, protein purification, VT - experimental design, mutagenesis, protein purification, spin labeling, reconstitution, data analysis, writing and editing manuscript.

